# Bioengineering a Miniaturized In Vitro 3D Myotube Contraction Monitoring Chip For Modelization of Muscular Dystrophies

**DOI:** 10.1101/2021.06.15.448543

**Authors:** Nicolas Rose, Surabhi Sonam, Thao Nguyen, Gianluca Grenci, Anne Bigot, Antoine Muchir, Benoît Ladoux, Fabien Le Grand, Léa Trichet

## Abstract

Quantification of skeletal muscle functional contraction is essential to assess the outcomes of therapeutic procedures for muscular disorders. Muscle three-dimensional “Organ-on-chip” models usually require a substantial amount of biological material, which is problematic in the context of limited patient sample. Here we developed a miniaturized 3D myotube culture chip with contraction monitoring capacity. Optimized micropatterned substrate design enabled to obtain high culture yields in tightly controlled microenvironments. Spontaneous contractions in myotubes derived from primary human myoblasts were observed. Analysis of nuclear morphology confirmed a similar organization between obtained myotubes and *in vivo* myofibers. *LMNA*-related Congenital Muscular Dystrophy (L-CMD) was modelled with successful development of mutant 3D myotubes displaying contractile dysfunction. This technology can thus be used to study contraction characteristics and evaluate how diseases affect muscle organization and force generation. Importantly, it requires significantly fewer starting materials than current systems, which should allow to substantially improve drug screening capability.

## 1. Introduction

Skeletal muscles constitute the most prevalent tissue in the human body, representing more than half the corporal mass. Force generation relies on the contractile features of the myofibers constituting the muscle tissue. Myofibers are long multinucleated cells containing contractile myofibrils organized in periodic segments. These segments, called sarcomeres (1), are composed of actin and myosin filaments parallel to the myofiber axis. Sliding of these myo-filaments along one another results in sarcomere shortening, and subsequent myofiber contraction (2).

Myofibers are terminally differentiated and cannot repair themselves in case of trauma. Yet, they host muscle stem cells (MuSCs) in niches between the myofiber membrane and the surrounding basal lamina (3). After an acute injury or a mechanical overload, MuSCs contribute to the muscle regeneration in a process tightly regulated by physical and chemical constraints of the niche. Return of skeletal muscle tissue to homeostasis following injury is classically evaluated by histology on cross-sections and force measurements as endpoints. As a complement to physiological symptoms analysis, there is a medical need to assess the muscle function in patients suffering muscular pathologies, for healthcare and research purposes (4). Indeed, muscle contraction is altered in the context of such rare dystrophies and quantification of skeletal muscle functional strength is essential to estimate the outcomes of therapeutic procedures.

The most common *in vitro* model used to study muscle is dish culture of myoblast monolayers, allowing studying early phases of myogenesis until myotube formation (5, 6), but when myotubes reach a certain level of maturation, contraction leads to detachment (7) from the 2D culture substrates. Therefore, investigating myotube contraction requires to adapt 3D tissue force measurement systems. Among them, methods based on micropillar substrates allow measuring myotube contractility (8–12). Fine-tuning of the pillar geometry allows to control the physical features of the forming tissue environment In parallel micro- and nanopatterned grooves with linear striations have been shown to promote cell alignment on 2D substrates (13, 14) and orientation of myocytes and cell fusion (15, 16). Patterned distribution of adhesive proteins also represents a powerful tool to spatially constrain living cells (17) in particular during myotube fusion and development (18). Light-induced molecular adsorption (LIMA) that consists in combining PLL-PEG polymers substrate passivation with UV local restoration of surface adhesion (19) can be used to homogeneously pattern surfaces having relief microfeatures, still this technique has not been yet implemented to skeletal muscle 3D organoids.

Powerful tools to perform therapeutical analysis (7, 20–21) already exist, but their important sizes require a high number of myoblasts to generate muscle micro-tissues. “Organ-on-chip” and “tissue-on-chip” technologies allow reducing the biological material prerequisite, by rescaling the dimensions of the organoids and expanding the number of replicates made from a constant initial sample size. Bioengineering research for such technology enabled to reduce the cell/construct ratio (22–25). In this work we propose a hydrogel free system to further diminish the size needed for a single tissue and increase the pillar density on the silicone chip. To do so, the geometrical features of the pillars and the adhesive proteins patterning technique were optimized to maximize the yields of 3D myotubes culture. This system enabled to monitor spontaneous contractions and study nuclei three-dimensional morphology of myotubes obtained from primary myoblasts as well as from healthy and L-CMD (LMNA-related Congenital Muscular Dystrophy) patients’ immortalized myoblasts. In our hands, the generation of single-myotube micro-tissues required 10 to 1000 less cells compared to currently available 3D skeletal muscle constructs.

## 2. Results

### 2.1. Downsized model for in vitro functional skeletal muscle culture

The goal of this study was to create a 3D myotube culture chip with contraction monitoring capacity for therapeutical research. To this aim, we developed a method that consists in culturing human myoblasts amplified from patients’ biopsies on microstructured substrates made in PDMS (Polydimethylsiloxane) (Fig. 1A). The pre-molded PDMS chip micro-substrates (10×10mm) (Fig. 1B) allow to culture micro-tissues between micropillar duos at a high density (each chip can easily be placed in a 24-well box well and can potentially bear 100 micro-tissues), providing numerous replicates while requiring relatively few myoblasts to be seeded (100k cells needed per chip). Pillar duos present in between the pillars either a pattern of 6 μm periodic striations, with a height of 3.5 μm (Fig. 1C), or no striation (controls). Different shapes, sizes and heights of the pillars were considered (Fig. 1B,D,E and table S2). Depending on their geometry, the pillars have different spring constant values (26), offering the possibility to adapt the resisting elastic force of the system to 3D myotube contraction.

**Fig. 1.**
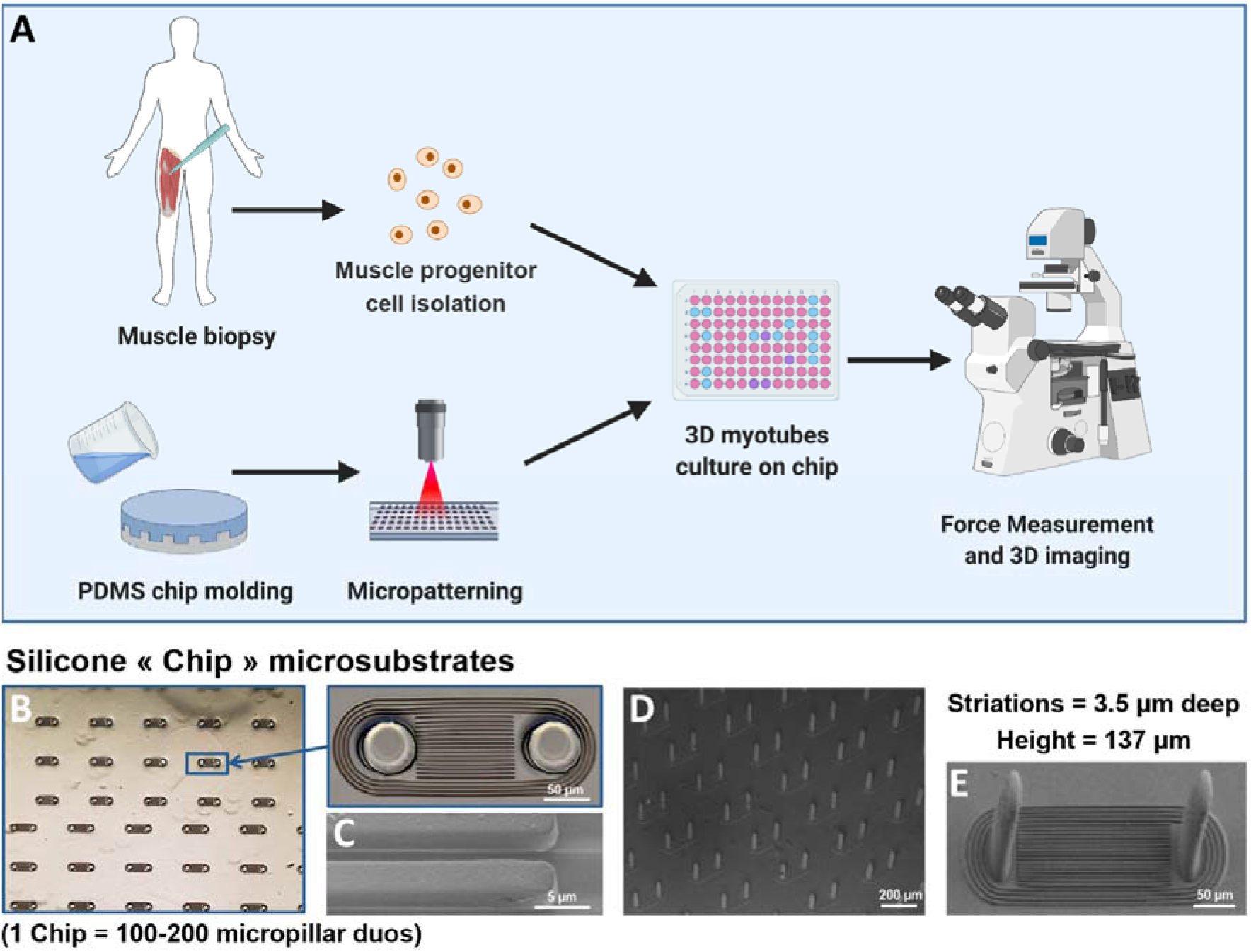
Methodology for developing an ex vivo system for micro-muscle contraction. **(A)** Schematic representation of the method, combining photopatterning technique with culture of patient muscle cells (PDMS=Polydimethylsiloxane) (Created with BioRender.com). **(B)** Transmitted light microscopic image of a chip’s topography, and observation of a pillar duo at a higher magnification. **(C,D,E)** Tilted scanning electronic microscopy images (30°) from few different pillar types. **(C)** striations, **(D)** pattern n°6 (table S2), **(E)** pattern n°2 (table S2).

### 2.2. Matrigel coating micropatterning

To gain in 3D myotube density, micropillar duos were placed side by side on the chip without recessed wells to receive the seeded cells. We used LIMA technique (Fig. 2A) to avoid myoblast monolayer formation over the entire chip surface and restrict myoblasts adhesion in the inter-pillar area. This technique is based on the “passivation” of the substrate by PLL-PEG polymers. While the poly(L-lysine) backbone interacts electrostatically with the substrate, the poly(ethylene glycol) side chains block proteins or cell adhesion (19). Combined to a photo-initiator, local exposure to UV degrades PLL-PEG, creating a region permissive to cell adhesion (Fig. 2B). The specificity of the Matrigel micropatterning was proved as cells would almost exclusively adhere to patterned areas from early proliferation phase to later stages of myotube maturation (Fig. 2C). Diverse technical parameters were optimized, such as the UV exposure (the required minimal dose being 1000 mJ/mm^2^ for optimal patterning) of PLL-PEG polymers (Fig. 2D). The developed protocol enables to obtain Matrigel-coated areas exclusively between the pillar duos (Fig. 2E), forcing the myoblasts to adhere in clusters and fuse into myotubes strictly in the axis of the pillars (Fig. 2F). Micropatterning-based clustering of cells appeared to be an essential element of culture success, since hydrogel embedment in both pure Matrigel or Matrigel/Collagen I mix, allowing branching between myotubes, displayed disastrous effects on 3D myotube culture yields (Fig. S2), whereas controls on 2D monolayer cultures confirmed promotion of myotube maturation within Matrigel embedment (Fig. S2A-E)(5). The system was kept gel-free with only silicone substrate and liquid medium to support 3D myotube culture.

**Fig. 2.**
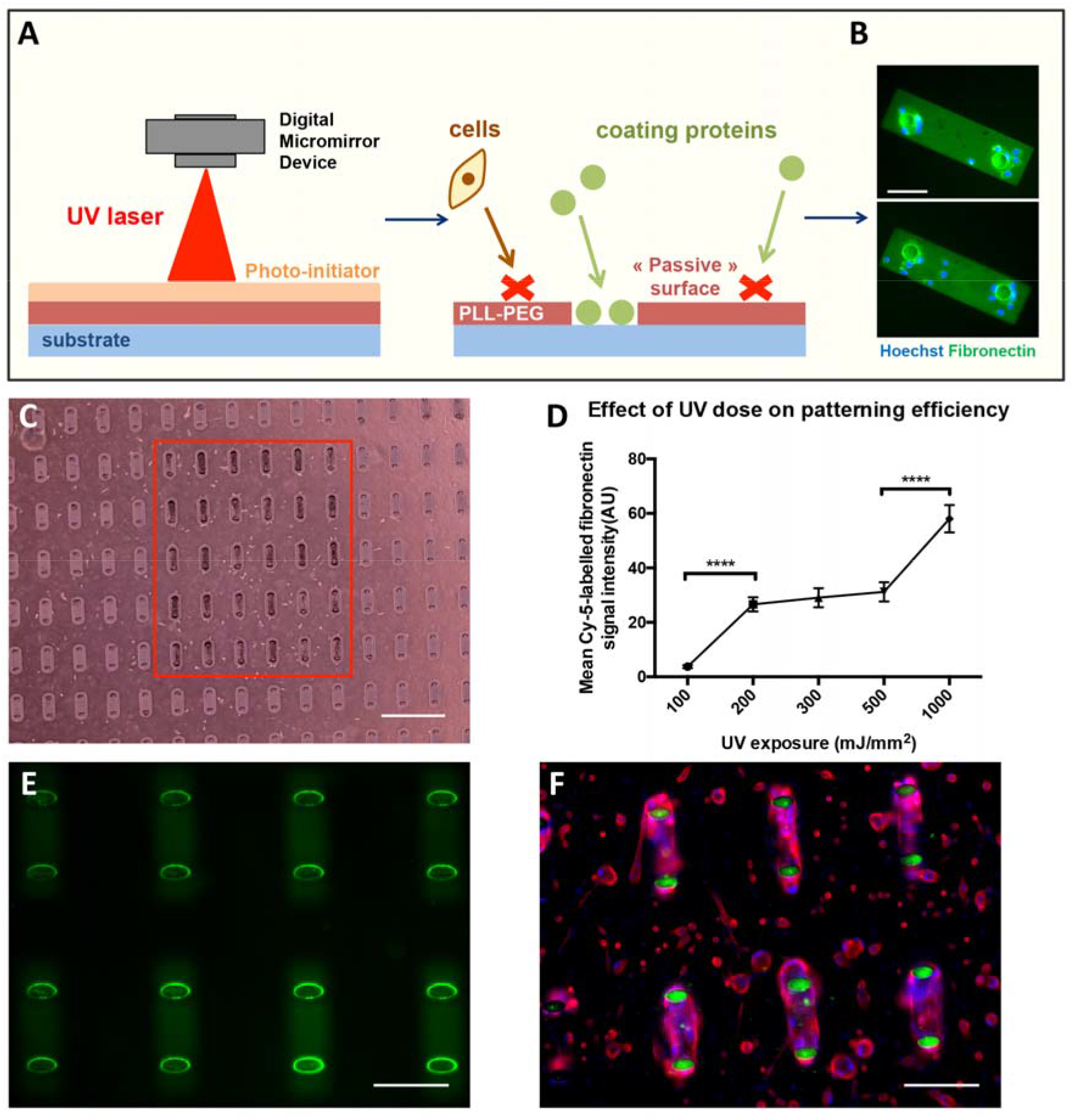
Micropatterning 3D myotubes with light induced molecular adsorption. **(A)** Schematic representation of LIMA technique. **(B)** Images of immunostained human immortalized fibroblasts nuclei (blue) plated on microsubstrates with Cy5-labeled fibronectin (green). Scale bar = 100 μm. **(C)** Transmitted light microscopic image of 30 microtissues obtained from human immortalized (8220) myoblasts seeded the day before, forming within patterned area (red rectangle). Scale bar = 500 μm. **(D)** Mean Cy-5 labelled fibronectin fluorescent signal intensity (± s.e.m.) measured after UV exposure along a spatial gradient. ****p<0.0001 (Unpaired t-test). **(E)** Image of micropillars patterned with Fibronectin-Cy5/Matrigel mix (FN-Cy5 = green). Scale bar = 200 μm. **(F)** Image of immunofluorescent staining of forming myotubes from human immortalized (8220) myoblasts after 7 days of differentiation on micropillars patterned with FN-Cy5/Matrigel mix (Myosin heavy chain (red), FN-Cy5 (green), nuclei (blue). Scale bar = 200 μm.

### 2.3. Optimization of chip geometrical settings

To facilitate myoblast alignment, 6 μm periodic striations, with a height of 3.5 μm, were sculpted in between the micropillar duos, as this pattern was already shown to produce the healthiest, aligned myoblasts when comparing to 3 and 12 μm periodicity size (15). This topographical modification drastically improved 3D myotube yield (Fig. 3A-C). We only obtained a success rate (namely, the ratio of observed myotubes in between pillar duos after 7 days of differentiation over total number of prepared pillar duos) of 6% for immortalized myoblasts-derived 3D myotube without striations, whereas a 35% success rate on striated ones was achieved. Results were collected from different pillar geometries (table S2: patterns n°3, 4, 7 and 8, N=4 for each pattern). This percentage was defined as the ratio of observed myotubes in between pillar duos after 7 days of differentiation over total number of prepared pillar duos. After fusion, the successfully formed myotubes embraced the pillars. Interestingly, we observed that during the maturation of the myotubes, their spontaneous contraction basically rips them of the surface inducing their lateral migration upward and leading to the bending of the two elastic pillars toward each other. We then analyzed the impact of pillar geometry to determine the optimal conditions of myotube formation. The 3D myotube obtention rate decreased linearly with increasing pillar major axis (fig. S1). Two pillar shapes, presenting elliptic shapes with axial ratios close to 1.7 (table S2: patterns n°5 and 6) provided optimized 3D myotube yields. After migration the microtissues remain suspended between these pillars at an average height of 75 μm (SD=11μm, N=30), independently of cell type as confirmed by observation on brightfield images and by scanning electron microscopy (SEM) (Fig. 3D). Success rates reached respectively 80 %, for a major axis of 54 μm, and 58% for a major axis of 44 μm, after 7 days of differentiation, and 63 % and 58% after 10 days of differentiation (Fig. 3E). The pillar deflection can also be easily observed from the top by transmitted light microscopy, allowing to estimate the generated force, calculated from elastic beam with ellipsoidal cross-section (Fig. 3F) (table S2: n°1, 2, 4, 5, 6). Using these optimized parameters each 3D myotube requires 1000 to 6667 seeded myoblasts, considering that 100k cells are plated to obtain 100 myotube/chip in optimal conditions, whereas 200k cells are plated to obtain 30 myotube/chip in the worst conditions, such as the most severe mutant phenotype.

**Fig. 3.**
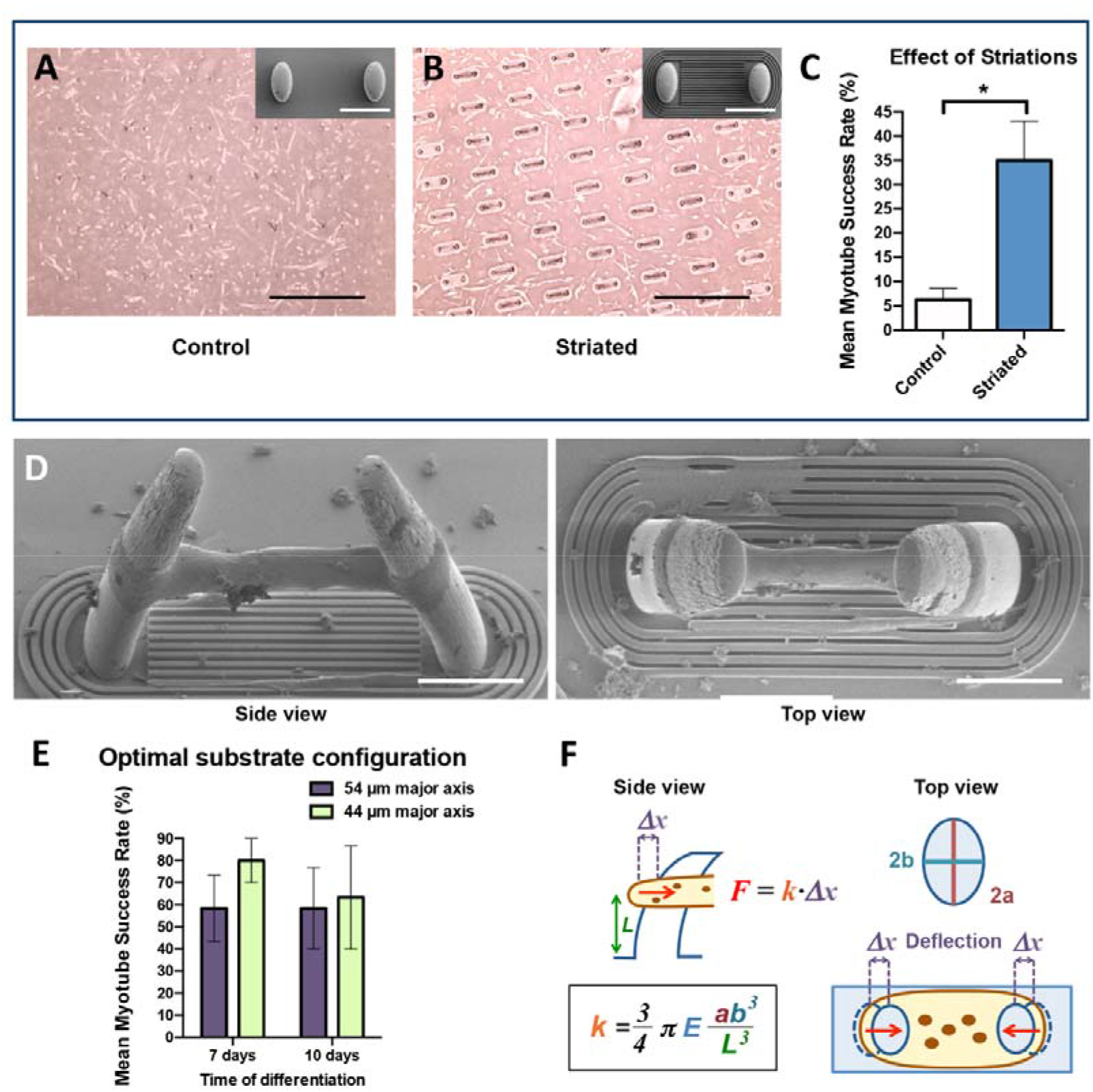
Optimization of substrate configuration for myotube culture efficiency. **(A,B)** Transmitted light microscopic images of myotubes obtained from human immortalized myoblasts (8220 and AB1190) after 7 days of differentiation (scale bars = 100 μm), and associated scanning electronic microscopy images (scale bars = 600 μm) of corresponding non striated (A) and striated (B) identical micropillar duos (patterns n°7 and n°4 - table S2). **(C)** Analysis of mean 3D myotube culture success rate ± s.e.m. obtained from human immortalized (8220 and AB1190) myoblasts after 7 days of differentiation on patterned micropillars with or without 6 μm periodic striations (pillar n°3, n°4, n°7 and n°9; see table S2). ± s.e.m. is represented, \**p*<0.05 (Wilcoxon matched-pairs signed rank test), N=4. **(D)** SEM images (top view and tilted view 45°) of a 3D myotube obtained from human immortalized (8220) myoblasts after 7 days of differentiation (pattern n°6 - table S2). Scale bars = 50 μm. **(E)** Analysis of mean myotube obtention success rate (± s.e.m.) from human immortalized (8220 and AB1190) myoblasts after 7 and 10 days of differentiation on patterned micropillars with the two most optimal substrate configurations (pillar n°5 in purple and n°6 in green. See table S2), N=2 (analysis was performed on at least 60 patterned pillar duos for each cell line). **(F)** Calculation of the bending force per pillar with ellipsoidal cross-section (F: generated force; k: Spring constant (μN/μm)(34); E: PDMS 1:10 Young’s modulus = 1.8 MPa; 2a: major axis; 2b: minor axis; L: vertical position of the microtissue on the pillars).

### 2.4. Functional analysis of 3D myotubes

Maturated 3D myotubes from primary human myoblasts cultured on micropillars successfully developed a sarcomeric network (Fig. 4A), which spreads spatially in a cylindrical fashion (Fig. 4B), according to the myotube shape, and expressing markers of the contractile apparatus (Fig. 4C and 4D) (e.g. titin and sarcomeric α-actinin respectively staining I-bands and Z-disks). The nuclei also aligned along the myotubes at different height levels (Fig. 4E), similarly to myonuclei observed on isolated myofibers. Following the formation of the contractile machinery, we observed input-free twitch contractions of our 3D myotubes (Movie S1) (fig. S3) (table S2: pattern n°6), which could be monitored by pillar deflection tracking (Fig. 4F-G). Unlike most preceding models using optogenetic or electrophysiological activations (24, 27), we observed contraction events of the myotubes without exogenous stimulation, after myofibril expression has been reached. These contractions were recorded on primary (Fig. 4F) (Movie S1) and immortalized (Fig. 4G) (Movie S2) myoblasts-derived 3D myotubes (with respectively 44 and 183 monitored contractions phenomena). Primary myoblasts-derived 3D myotubes systematically displayed a repetitive and stereotyped pattern of contractions within approximately 1.5 second, with a stability of the force value normalized by the number of nuclei for each myotube (Fig. 4H), notably the value corresponding to the peak (Fig. 4I). We first observed a fast increase of the force until the peak of the contraction (after 0.5 second), followed by a progressive force decrease (lasting 0.5 to 1 second). Immortalized myoblasts-derived 3D myotubes displayed a similar pattern of contraction, but with a reduced intensity (1.14 +/− 0.04 μN instead of 2.44 +/− 0.06 μN) in a reduced time frame (0.43 +/− 0.01 second instead of 1.03 +/7−0.03 second) (Fig. 4H-I). The frequency is four times superior, with values respectively of 1.11 Hz for the immortalized myoblasts (Fig. 4G) and 0.27 Hz for the primary myoblasts (Fig. 4F). Furthermore, immortalized myoblasts-derived 3D myotubes displayed a strong variability of spontaneous contraction intensity in comparison to primary myoblasts-derived 3D myotubes (with respectively 44.77% and 9.28% variation coefficients).

**Fig. 4.**
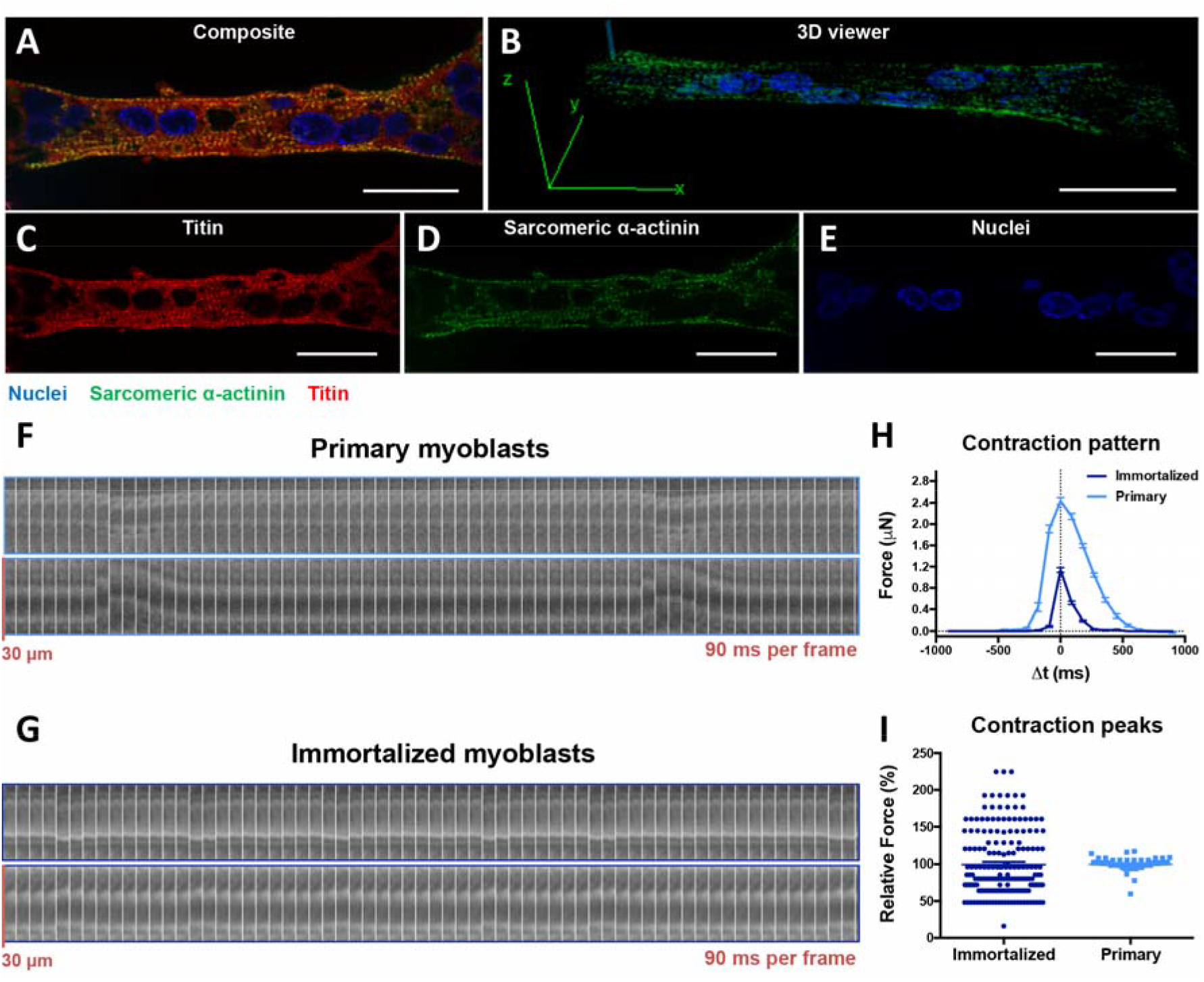
Monitoring of spontaneous contraction generated by mature 3D myotubes. **(A,B,C,D,E)** Representative images of immunofluorescent stained 3D myotubes obtained from human primary (AB678C53Q) myoblasts after 12 days of differentiation: confocal section (A,C,D,E) and three-dimensional view (B) (Sarcomeric α-actinin (green), Hoechst (blue), Titin (red), scale bars = 25 μm). **(F,G)** Transmitted light kymographs of pillar deflections over time with 3D myotubes from primary (AB678C53Q) (F) and immortalized (8220) (G) myoblasts after 9 days of differentiation. **(H)** Mean relative contraction force over time (Normalized with nuclei number: Each force was multiplied by this nuclei ratio = (100 / nuclei number) to be standardized based on a 100 nuclei tissue; considered height: 75 μm) ± s.e.m. generated by 3D myotubes from human primary (AB678C53Q) (light blue) and immortalized (8220) (dark blue) myoblasts after 9 days of differentiation. (pillar n°6. See table S2) (primary: N=2; 44 contractions (dots) observed, immortalized: N=3; 183 contractions (dots) observed). **(I)** Relative forces of contraction peaks (normalized with mean peak contraction force) ± s.e.m. generated by 3D myotubes from human primary (AB678C53Q) (light blue) and immortalized (8220) (dark blue) myoblasts after 9 days of differentiation (primary: N=2; 44 contractions (dots) observed, immortalized: N=3; 183 contractions (dots) observed).

### 2.5. Morphological impact of 3D culture on myonuclei

Muscle disorders such as laminopathies are known to affect myonuclei structure, which have also been found to display variation in shape when cultured in 3D rather than 2D in previous skeletal muscle organoids (21). To investigate the morphological impact of 3D culture on nuclei shape in comparison to 2D culture, we analyzed the 3D myotube organization at different length scale. We measured each nucleus projection area slice by slice on a vertical z axis, as well as their 3D features (Fig. 5A-D). Representation of mean normalized area (with regard to Maximal Slice Area – MSA) as a function of the distance from the nucleus center gives an indication of the 3D shape, with steeper slopes corresponding to flatter nuclei. The distributions of nuclei slice areas showed that nuclei within 3D myotubes presented largely reduced flatness compared to nuclei from 2D monolayers (Fig. 5E). However, nuclei from 3D myotubes conserved a similar ellipticity (Fig. 5F), showing that the nuclear width and length evolved in the same proportions. In comparison to nuclei from 2D monolayers, nuclei from 3D myotubes had heights twice superior (respectively 3.2 μm and 7 μm) (Fig. 5G). In addition, 3D nuclei also exhibited decreased volume (Fig. 5H) compared to 2D nuclei (respectively 229 μm^3^ and 285 μm^3^).

**Fig 5.**
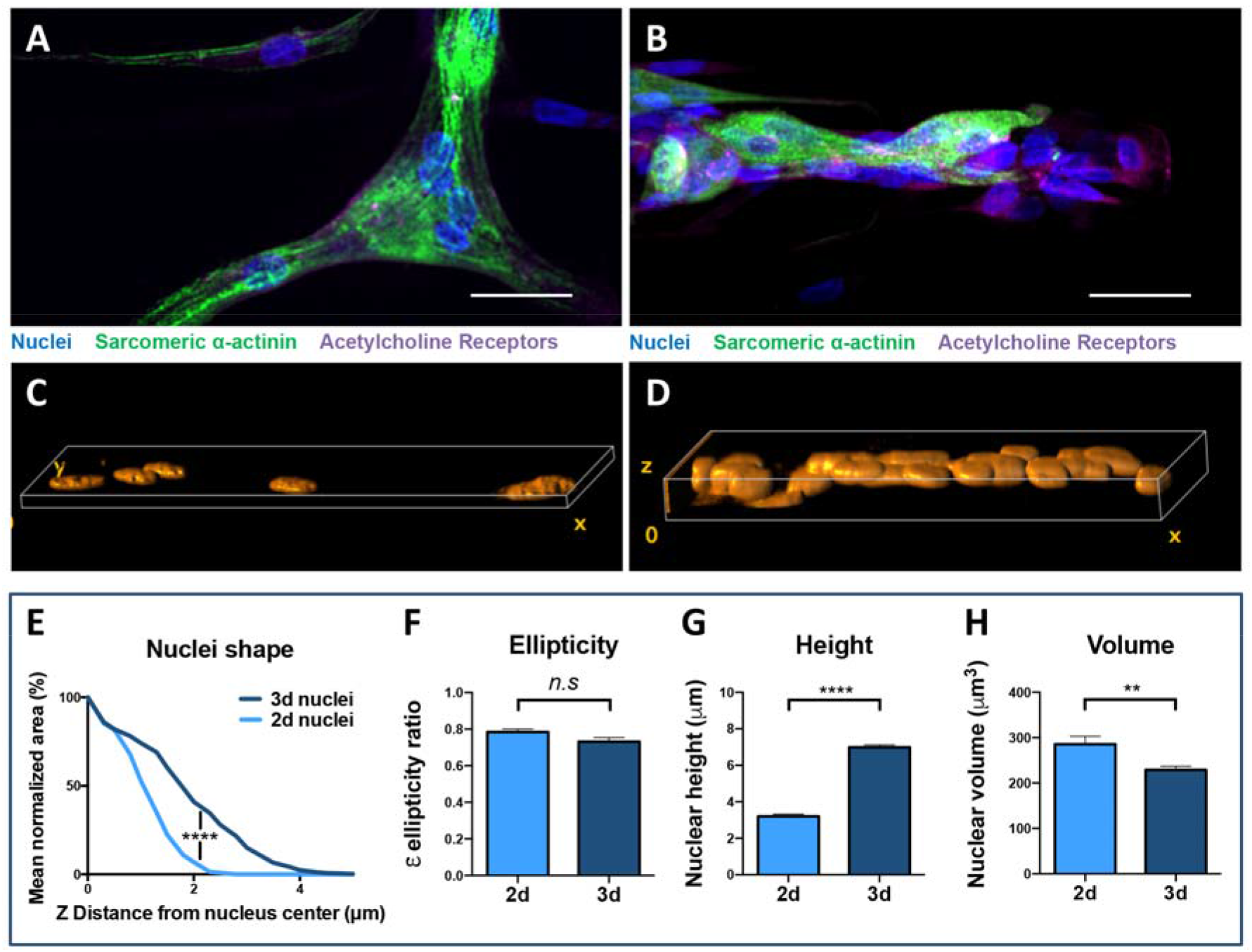
Shape and dimensions of nuclei within 2D monolayer and 3D myotubes. **(A,B)** Representative (top view) images of an immunofluorescent stained monolayer of myotube (A) and 3D myotube (B) obtained from human primary (AB678C53Q) myoblasts after 7 days of differentiation (Sarcomeric α-actinin (green), Hoechst (blue), Acetylcholine receptors (magenta) scale bars = 25 μm). **(C,D)** Volume viewer of Hoechst-stained nuclei from tilted representative images of myotubes monolayer (C) and 3D myotube (D) obtained from human primary myoblasts after 7 days of differentiation (rotations : x=105; y=15; x=0). **(E)** Distributions of mean normalized areas calculated in 2D myotubes monolayers and 3D myotubes obtained from human primary (AB678C53Q) myoblasts after 7 days of differentiation. \*\*\*\**p*<0.0001 (Wilcoxon matched-pairs signed rank test), 2D = 35 nuclei; 3D = 69 nuclei. (F) Mean nuclear (xy) ellipticity ± s.e.m. in 2D myotubes monolayer and 3D myotubes obtained from human primary myoblasts after 7 days of differentiation. *n.s* not significant (Mann-Whitney unpaired test), 2D = 35 nuclei; 3D = 69 nuclei. **(G)** Mean nuclear height ± s.e.m. in 2D myotubes monolayer and 3D myotubes obtained from human primary myoblasts after 7 days of differentiation. \*\*\*\**p*<0.0001 (Unpaired t-test), 2D = 35 nuclei; 3D = 69 nuclei. (H) Mean nuclear volume (± s.e.m.) in 2D myotubes monolayer and 3D myotubes obtained from human primary myoblasts after 7 days of differentiation. \*\**p*<0.01 (Mann-Whitney unpaired test), 2D = 35 nuclei; 3D = 69 nuclei. (pillar n°6. See table S2).

### 2.6. Expression of pathological nuclear phenotypes in patient 3D myotubes

We then used our device to investigate the mechanical differences between 3D myotubes from healthy (Fig. 6A) and L-CMD (Fig. 6B) patients immortalized myoblasts. As observed in 3D reconstitutions from confocal imaging (Fig. 6C-D), nuclei with *LMNA* mutation displayed morphological differences compared to control nuclei. First, *LMNA* mutation was associated to a significant increase of nuclear flatness (Fig. 6E-F), in comparison to control nuclei. Furthermore, while maintaining a constant nuclear ellipticity (Fig. 6G), 3D myotubes issued from L-CMD patients also differed from control myotubes by presenting longer (respectively 14.5 μm and 11.4 μm) and wider (respectively 9.6 μm and 7.1 μm) nuclei (Fig. 6H-I). Confirming the relative flatness of LMNA mutated nuclei, the height (Fig. 6J) was slightly but significantly reduced compared to control nuclei (respectively 5.2 μm and 5.8 μm), despite the larger volume (Fig. 6K) of mutant nuclei (211 μm^3^ against 162 μm^3^ in control nuclei). Altogether, these results showed that once put in our system, patient-derived myoblasts can generate 3D myotube reproducing typical L-CMD pathological phenotype which have previously been reported (21, 28).

**Fig. 6.**
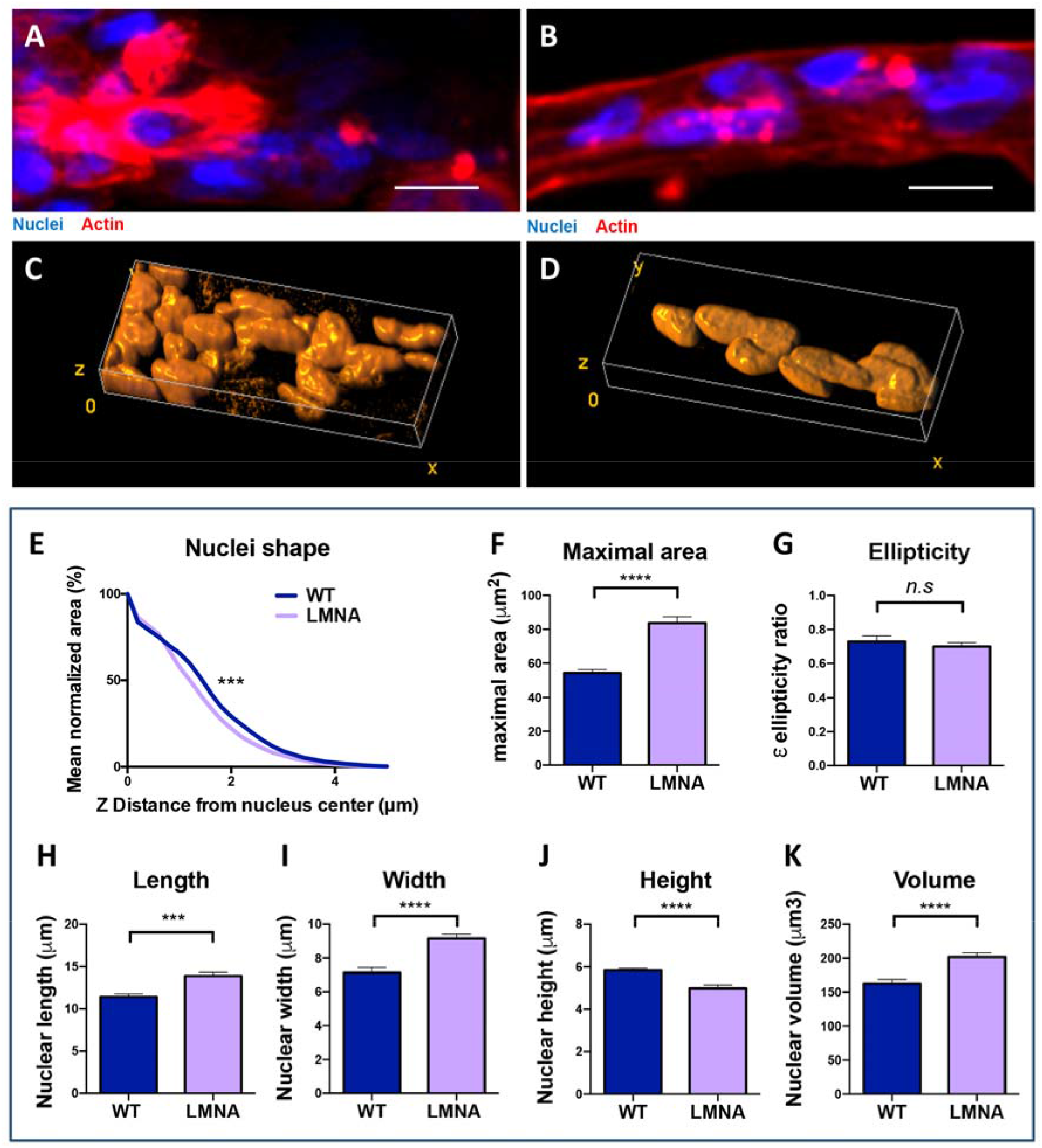
Effect of a LMNA gene mutation on nuclei morphology in patient 3D myotubes. **(A,B)** Representative (top view) images of immunofluorescent stained 3D myotubes obtained from control (A) and L-CMD (B) patients myoblasts (respectively 8220 and EMD1365) after 7 days of differentiation (Hoechst (blue), phalloidin (red) scale bars = 25 μm, both images are later analyzed in C and D). **(C,D)** Volume viewer of Hoechst-stained nuclei from tilted representative images of 3D myotubes from control (C) and L-CMD (D) patients myoblasts after 7 days of differentiation (rotations : x=150; y=15; x=10). **(E)** Distributions of mean normalized areas calculated in 3D myotubes from control (8220) and L-CMD (EMD1365) patients’ myoblasts after 7 days of differentiation. ***p<0.001 (Wilcoxon matched-pairs signed rank test), WT = 57 nuclei; LMNA = 85 nuclei. **(F)** Mean area at the central slice of the nuclei ± s.e.m. ****p<0.0001 (Mann-Whitney unpaired test). **(G)** Mean nuclear (xy) ellipticity ± s.e.m. in 3D myotubes from control (8220) and L-CMD (EMD1365) patients’ myoblasts after 7 days of differentiation. n.s not significant (Unpaired t test), WT = 57 nuclei; LMNA = 85 nuclei. **(H,I,J,K)** Mean nuclear length (H), width (I), height (J) and volume (K) ± s.e.m. in 3D myotubes from control (8220) and L-CMD (EMD1365) patients myoblasts after 7 days of differentiation. ***p<0.001; ****p<0.0001 (Mann-Whitney unpaired test), WT = 57 nuclei; LMNA = 85 nuclei. (pillar n°6. See table S2).

### 2.7. Contractile dysfunction in L-CMD patients 3D myotubes

To assess the functional features of pathological constructs, we monitored pillar deflections caused by the contractile activity of 3D myotubes issued from healthy donors (Fig. 7A) (Movie S2) and L-CMD patients (Fig. 7B) (Movie S3) immortalized myoblasts. We observed smaller deflections of the pillars in 3D myotubes carrying *LMNA* mutation (Fig. 7C) as compared to control myotubes. Unlike in control myotubes, spontaneous twitch contractions could not be observed in *LMNA* mutant myotubes. These amplitudes were also both significantly higher in comparison to inactivated 3D myotubes (pre-treated with 20 μM Acetylcholine) (Fig. 7C). Moreover, the difference in absolute pillar deflection amplitude could not be observed when normalized with nuclei number (Fig. 7D). Indeed, LMNA mutant myotubes contained less nuclei than the control myotubes (with respective means of 62.7 and 80.0 nuclei per myotube, data not shown). This suggests that these lower amplitudes and associated weaker forces are related to a reduction in the efficiency of muscle cell fusion and subsequent myotube formation in LMNA-mutant context.

**Fig. 7.**
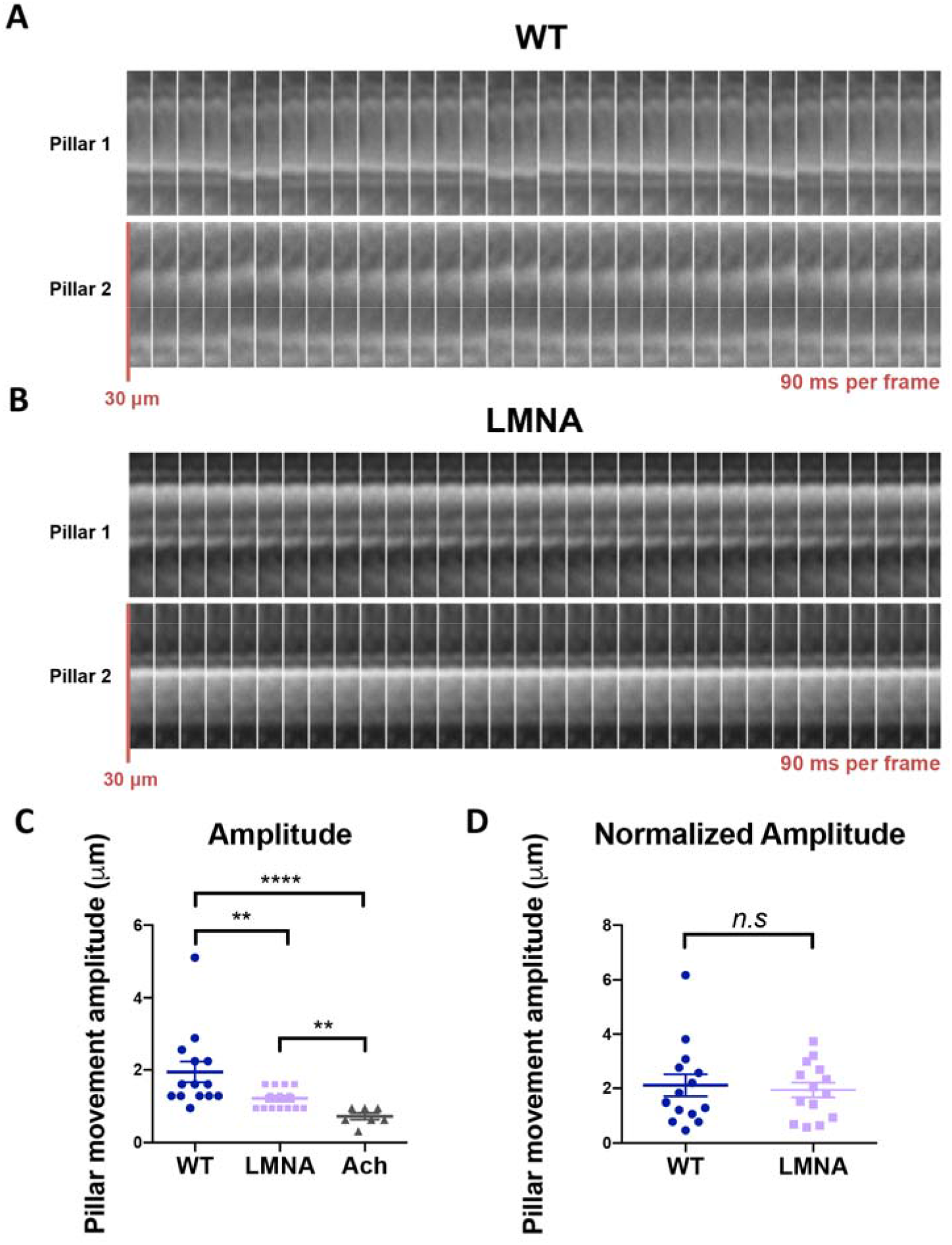
Effect of a LMNA gene mutation on contractile activity of 3D myotubes obtained from patients’ myoblasts. **(A,B)** Transmitted light kymographs of pillars deflection over time with 3D myotubes from control (8220) (A) and L-CMD (EMD1365) (B) patients immortalized myoblasts after 9 days of differentiation. **(C)** Mean maximal amplitude of movement ± s.e.m. (measured on 9 seconds films) of pillars with 3D myotubes from control (8220) or L-CMD (EMD1365) patients’ myoblasts and 3D myotubes inactivated with 20 μM Acetylcholine, after 9 days of differentiation. **p<0.01; ****p<0.0001 (Mann-Whitney unpaired test), N=14 (and 7 Acetylcholine-inactivated myotubes). **(D)** Mean maximal amplitude of movement (Normalized with nuclei number: Each force was multiplied by this nuclei ratio = (70 / nuclei number) to be standardized based on a 70 nuclei tissue) ± s.e.m. (measured on 9 seconds films) of pillars with 3D myotubes from control (8220) and L-CMD (EMD1365) patients’ myoblasts after 9 days of differentiation. *n.s* = not significant (Mann-Whitney unpaired test), N=14.

In conclusion, the micro-muscle chip developed here can accurately measure differences in force generation by constructs generated from muscle cells expanded from patients with genetic muscular diseases.

## 3. Discussion

We developed a tissue-on-chip culture device allowing the assessment of forces generated by 3D myotubes from healthy donors or from patients suffering muscular disorders. The system relies on a combination of optimized geometrical features of the substrate, namely pillar duos linked together by micropatterned linear striations, with adhesive proteins micropatterning mediated by light-induced molecular adsorption. Fine tuning of each parameter contributes to increasing skeletal muscle organoids yield, and the underlying principles allow for further downsizing of the system. Thereby, this technology offers perspectives for implementation in myology research and organ-on-chip bioengineering for medical studies. This model might offer less versatility than other muscle constructs (21, 25, 27), which recreate muscle-like beams of fiber bundles, and allow co-culture of vascular (21) and neuronal (25, 27) tissues. Also, the presence of residual myocytes between patterned areas (Fig. 2F) could eventually bias bulk RNA and protein based molecular biology approaches performed with other models (20, 24). However, we propose a culture device with an unprecedented cells-to-construct ratio designed to produce a great number of technical replicates with relatively few cells. Each 3D myotube requires 1 000 to 6 667 seeded myoblasts depending on the cell type (in particular, primary or immortalized, control or mutant). For comparison purposes, 3D Artificial Muscles (21) requires one million of cells per construct (×10^3^ more), MyoTACTIC (27) uses gel suspension with 15 millions cell/mL, NMJ chip (25) have an estimated biological cost of 84 000 cells per tissue and human skeletal micro muscles (24) need 30 000 cells per tissue (×10^1^ more). Downsizing of the system could still be optimized by increasing pillar density to maximize 3D myotube culture yield.

The system has been optimized with diverse cell lines, primary or immortalized myoblasts, obtained from different muscle groups and from various patients with diverging ages and phenotypes (table S1). 3D myotubes were obtained for the two healthy myogenic immortalized lines. There is a strong reproducibility between experiments performed with the same cell line, which ensures the reliability of this platform when used to study the effect of a therapeutical drug on a patient cell line. Nevertheless, there is an unequal yield (increasing with duration of culture) depending on the cell line (data not shown), most likely due to uneven adhesion capacity, passive resting tension and sarcomere-generated contraction forces. Consequently, the chip configuration should ideally be fine-tuned for each myogenic lineage, in particular to control the physical parameters such as the stiffness constraints applied to the myotubes. In the present study 3D myotube culture was optimal using the two elliptic pillar shapes presenting the lowest major axis values (Fig. 3E). Counterintuitively, the tissue stability decreased linearly with increasing pillar major axis (fig. S1), suggesting that higher adhesion surfaces did not prevent myotube detachment. On the contrary, the wider myotubes seem to detach more easily, possibly due to increased contractile material.

Our system does not require the use of a hydrogel scaffold and rather relies on the production by the cells of their own matrix along the tissular formation process. Upward migration along the pillars is induced by spontaneous contraction of the tissues. Abstaining from any preexisting hydrogel allows proper pre-alignment of the seeded myoblasts in the substrate micropatterned striations, which is a major factor to ensure success of myotube formation and enables to induce mono-cylindrical shaping of the sarcomeric network. The formed myotube bundles are more compact with straighter orientation. Indeed, the alignment along the axis joining the pillars takes place in a steadier manner than in models using gel scaffolds (22–24). This is beneficial since variability in myotube density and orientation within the gel construct might create biases of force measurement.

Spontaneous twitch contractions in 3D myotubes derived from human myoblasts were observed with a high degree of reproducibility, highlighting the relevance of this technology. The specific forces of the spontaneous contractions here observed are in the order of few mN/mm^2^, which is comparable to the specific forces of twitch and tetanic contractions measured in other muscle microtissues (20, 24, 22, 27). This value is particularly interesting due to the spontaneous nature of the observed contractions, as opposed to the activated contractions reported in the literature. Contractions of primary tissues obtained with other reported protocols are induced by optogenetic (24, 27) or electrophysiological (20) stimulations. The observation of spontaneous contractions in our system highlights its relevance to study contractions in a clinical setting, as cells can be used directly from the patient’ biopsies. Still, spontaneous contractions were not systematically observed in every matured tissue. Use of specific drugs (Caffein, Calmodulin, Nocodazol, Rapamycin, Bosutinib, Reparixin, etc…), ions (Calcium, Potassium, etc…), and other molecular supports such as ATP (25) that are known to induce contractions, could be used to increase steadiness of obtained results. spontaneous contractions were also observed in immortalized myoblasts-derived 3D myotubes (Fig. 4) and presented different contraction characteristics when compared to primary myoblasts-derived 3D myotubes. The observed reduced and more frequent contractions could be explained by desynchronized activation of the different myofibrils within the 3D myotubes, leading to a fibrillatory pattern. Contractions generated by immortalized myoblasts-derived 3D myotubes also displayed important variability (Fig. 4I), which seems to indicate a higher relevance of the system for primary myoblasts contraction analysis in comparison to immortalized cells, and further highlights the interest of this miniaturized culture system to harness patients’ biopsies.

Our reduced-scale model allows to precisely image subcellular structure like nuclei, sarcomeric network and many others. As expected, in absence of direct physical constraints from the substrate, 3D myotube nuclei substantially lost their 2D flatness (Fig. 5). The nuclei kept a length superior to their width and height, which is most likely due to the longitudinal mechanical forces exerted within the myotube by the cytoskeleton. This nuclear phenotype could also be reproduced previously by exerting external force on gel-borne nuclei (28). The overall volume of 2D monolayer nuclei was found to be more important than the one of 3D myotube, confirming the relative morphological aberrance caused by 2D culture and proving the need for three-dimensional systems of culture. The shape of 3D myotube nuclei is closer to the in vivo conformation of myonuclei (29, 30), which makes our culture chip a relevant tool for the analysis of nuclei-linked pathology, such as laminopathies.

The importance of the lamin envelope for nuclei morphological integrity is well known (31). Mutations in *LMNA* gene, encoding Lamin A and Lamin C proteins, are leading to striated muscle laminopathies, among which L-CMD, is the most severe type (32). The mutated lamins A/C are causing nuclear defects such as chromatin protrusion and nuclear envelope rupture (33). The ΔK32-P1 mutant myoblast lineage (28) we cultured into 3D myotube expressed the same phenotype that has been previously observed in preceding 3D skeletal muscle platforms (21). *LMNA* mutated nuclei showed similar morphological aberrations such as increased flatness and volume. This proves that our 3D myotube culture chip is a reliable system to model morphological abnormalities of muscular disorders. In addition, *LMNA* mutant 3D myotube displayed reduced absolute contractile activities compared to control 3D myotubes (Fig. 7C). However, this difference could not be observed after normalization with the tissue size, assessed based on total nuclei number, suggesting that this contractile dysfunction is due to abnormal tissue formation and could be solved by improving early myogenesis of LMNA mutant tissue. We anticipate that our device can thus be implemented as a readout for therapeutic drug screening for myopathies.

## 4. Materials & methods

### 4.1. Silicon wafer production and silanization

Silicon wafers were provided by the Mechanobiology Institute (National University of Singapore). The pillars and the grooves were produced in a two-step etching procedure. Briefly, silicon oxide wafers (300 nm thick, thermally grown) were initially patterned with the layout comprising of micro-grooves and the opening for the pillars. Then, in the first step of dry silicon etching the micro-grooves were protected by a patterned layer of photoresist, thus only the opening left to produce the pillars were etched down. A boshlike process was used in an ICP reactor (SI500 RIE, Sentech, Germany) to etch the pillars cavities with vertical profiles. In the second etching step, the photoresist coating was removed and a second step of etching (same as before, but for shorter time) was then performed resulting in the fabrication of the micro-grooves together with reaching the final depth for the pillars cavities. Prior to PDMS casting, the silicon molds were coated with an anti-sticking layer of silane (1H,1H,2H,2H-perfluorooctyl-trichlorosilane 97%): after activation of the surface with O_2_ plasma, the mold is immediately placed in a vacuum jar, a drop of the silane is added in a small container and the jar is evacuated for 15 min and left for 2 h for silane vapor deposition.

### 4.2. Polydimethylsiloxane chip molding

PDMS Sylgard 184 silicone elastomer base was mixed with curing agent in a 1:10 ratio (Dow Corning, ref 01673921). The mixture was poured on the silicon wafer, degased (with vacuum) and heated at 80°C for 2 h. PDMS was then peeled-off from the silicon wafer and cut into individual chips.

### 4.3. Matrigel micropatterning

Chips were hydrophilized by 5 min exposure to plasma cleaning. PLL-g-PEG (SuSoS Technology) was added to the chip surface for 1 h and rinsed three times with Phosphate-Buffered Saline (PBS). Photoactivable reagent (Alvéole’s Classic PLPP 14.5 mg/mL) was added on the chip and specific areas were exposed with 1200 mJ/mm^2^, using PRIMO micropatterning device (Alvéole) on 20X objective, with Leonardo photopatterning software (Alvéole). Following the photopatterning step, photoactivable reagent was vigorously rinsed away with PBS at least 5 times. Matrigel matrix (Corning ref 354230) solution (1 mg/mL in DMEM) with 1:50 Cy5 dye-labeled fibronectin (conjugated in house using dialysis kit from Life technologies) was added on the chip and incubated for 1 h at room temperature. Matrigel solution was gently rinsed once with H_2_O, and the patterned chips kept in sterile water at 4°C, plated in a multiwell-plate protected from the light.

### 4.4. Myoblasts lines and cell culture

Skeletal myoblasts cell lines were provided by the Myoline platform from the Institut de Myologie, Paris. The features of patients and muscles of origin are described in the table S1. Cells were amplified in proliferation medium composed of DMEM (high glucose, GlutaMAX(TM), ThermoFischer scientific) with 16% 199 medium (Medium 199, GlutaMAX™ Supplement, ThermoFischer scientific), supplemented with 20% Fetal Bovine Serum (Gibco ref 15750-037), 5 ng/ml human epithelial growth factor (Life Technologies), 0.5 ng/ml bFGF (Life Technologies), 0.2 mM dexamethasone (Sigma-Aldrich), 25 μg/ml fetuin (Life Technologies), 5 μg/ml insulin (Life Technologies) and 0.5 μg/mL Gentamicine (Gibco). Cells were cultured directly in plastic dishes and the passages were performed with 0.05% Trypsine-EDTA (1X) (Gibco). Human primary myoblasts were used within less than 30 divisions.

### 4.5. 3D myotube generation

The chips were rinsed with sterile PBS, plated in new sterile multiwell plates and immersed with proliferation medium. Myoblasts were plated (100k to 200k cells per chip depending on cell type - primary or immortalized, control or mutant, etc…) directly above the chip in a 0.5 to 1 mL proliferation medium drop. The chips were then spinned at 300g for 5 min, and left at 37°C for 1 h. The medium covering the chips was then gently changed (while keeping the chip immersed) 3 to 4 times until the off-pattern cells would detach. The chip was kept in proliferating conditions overnight until a switch to differentiation medium (DMEM high glucose, GlutaMAX (TM) + 0.1 mg/mL Gentamicine), by removing 1mL from the culture well and adding 1mL of new medium (repeated 4 to 5 times). The chips were left in differentiation conditions for at least 6 days (partial medium change can be done every 2 days), and 0.4 μm/mL recombinant rat Agrin (R&D systems) in differentiation medium was then added to the 3D myotubes. Fresh differentiation medium (1 mL) was gently added before contraction monitoring (performed after 7 to 14 days of differentiation).

### 4.6. Fixation and immunostaining

Samples were fixed 10 min with 4% Paraformaldehyde aqueous solution (Electronic Microscopy Sciences), blocked with 4% Bovine Serum Albumin (BSA) and permeabilized with 0.1% TritonX-100. The following primary antibodies (diluted in 4% BSA) were used for immunostaining: mouse IgG2b anti-Myosin Heavy Chain Type I MYH7 (DSHB, 1:50), rabbit monoclonal anti-Titin (DSHB 9D10-s, 1:500), mouse IgG1 anti-Sarcomeric Alpha Actinin (Abcam 9465, 1:500), rat monoclonal anti-Nicotinic Acetylcholine Receptor (Sigma-Aldrich, 1:800). The following secondary antibodies (diluted in PBS) were used for immunostaining: goat Alexa-Fluor-488/546/647-conjugated anti-mouse IgG1/IgG2b, anti-rabbit and anti-rat (Life Technologies, 1:500).

Actin was stained with Phalloidin-tetramethylrhodamine conjugate (Santa-Cruz 362065, 1:500) and nuclei with Hoechst 33342 (Life technologies H3570, 1:10,000). Chips were mounted upside down on glass coverslip with Dako Fluorescence Mounting medium (ref 53023).

For scanning electron microscopy preparation, samples were fixed first in a 4% paraformaldehyde solution and second using glutaraldehyde in a cacodylate/saccharose buffer for 1 h at 4 °C. After washing in a cacodylate/saccharose buffer the samples were dehydrated through successive ethanol baths with increasing concentrations and subsequently dried at the carbon dioxide critical point.

### 4.7. Microscopy imaging

Transmitted light videomicroscopy for contraction monitoring was performed at 37°C and 5% CO2 on Nikon eclipse Ti microscope with coolsnap HQ2 camera and 20X objective. Fluorescent imaging was acquired with EVOS FL Cell Imaging System microscope (Life Technologies). Contraction force was calculated using Saez et al. 2007 formula (34). Confocal and 3D imaging was acquired with Nikon Ti2 microscope equipped with a motorized stage and a Yokogawa CSU-W1 spinning disk head coupled with a Prime 95 sCMOS camera (Photometrics). Scanning electron microscopy was performed with Hitachi SU-70 High vacuum FESEM 1 kV or 0.5 kV (high magnification) without coating.

### 4.8. Images and statistical analysis

Images were acquired using Metamorph software and further treated with ImageJ/FIJI. Nuclear volumes were calculated by addition of all individually measured xy areas and multiplication with the confocal Z-step (zy or xz areas and ellipticities were not considered). All graphs and statistical analyses were performed with Prism software (tests and p-values are described in the legend of each figure).

## Acknowledgements

The authors would like to thank Sylvie Hénon and Vincent Gache for fruitful discussion. Penney Gilbert is kindly acknowledged for careful reading of the manuscript Joseph d’Alessandro and Lakhmi Balasubramaniam are kindly acknowledged for their help with substrate preparation. The authors thank David Montero and the Institut des Matériaux de Paris Centre (IMPC FR2482) for servicing FEGSEM & EDX instrumentation. Cell lines were obtained thanks to the Myoline platform from the Institut de Myologie, Paris and to the Biobank support of the London MRC Neuromuscular Translational Research Centre and the Biomedical Research Centre of Great Ormond Street Hospital for Children in London. This work was supported by grants from the Agence Nationale de la Recherche (ANR-19-CE13-0016 MyoFuse), the Mechanobiology Institute (to G.G.), the LabEx “Who Am I?” #ANR-11-LABX-0071, the Université de Paris IdEx #ANR-18-IDEX-0001, the AFM/Téléthon and C’Nano program of the Région Ile-de-France. N.R. was supported by a French Ministère de la Recherche fellowship, Interfaces pour le vivant doctoral program from Sorbonne Université.

## Supplementary data

**Fig. S1.**
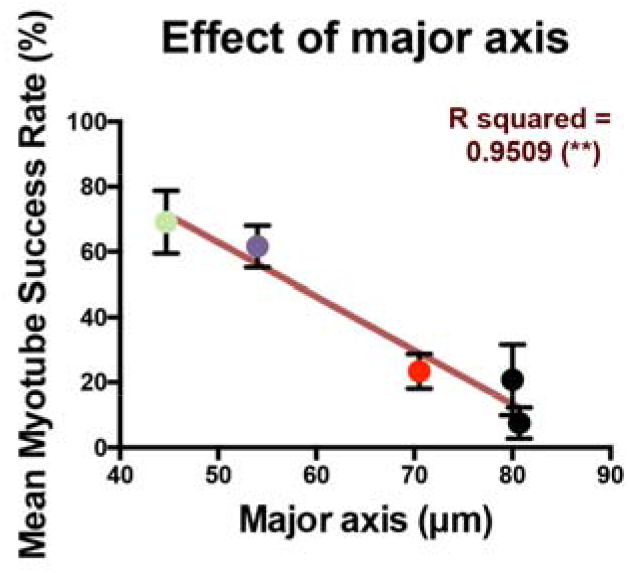
Effect of pillar major axis on culture yield. Analysis of 3D myotube obtention success rate ± s.e.m. obtained from human immortalized (8220 and AB1190) myoblasts after 7 days of differentiation on patterned micropillars with various major axis (pillar n°1, 2, 4, 5, 6 with 140 μm and 190 μm approximate interpillar distance. See table S2). Linear regression (red line) and Pearson correlation test (\*\**p*<0.01) and coefficient (R squared = 0.9509), N=4.

**Fig. S2.**
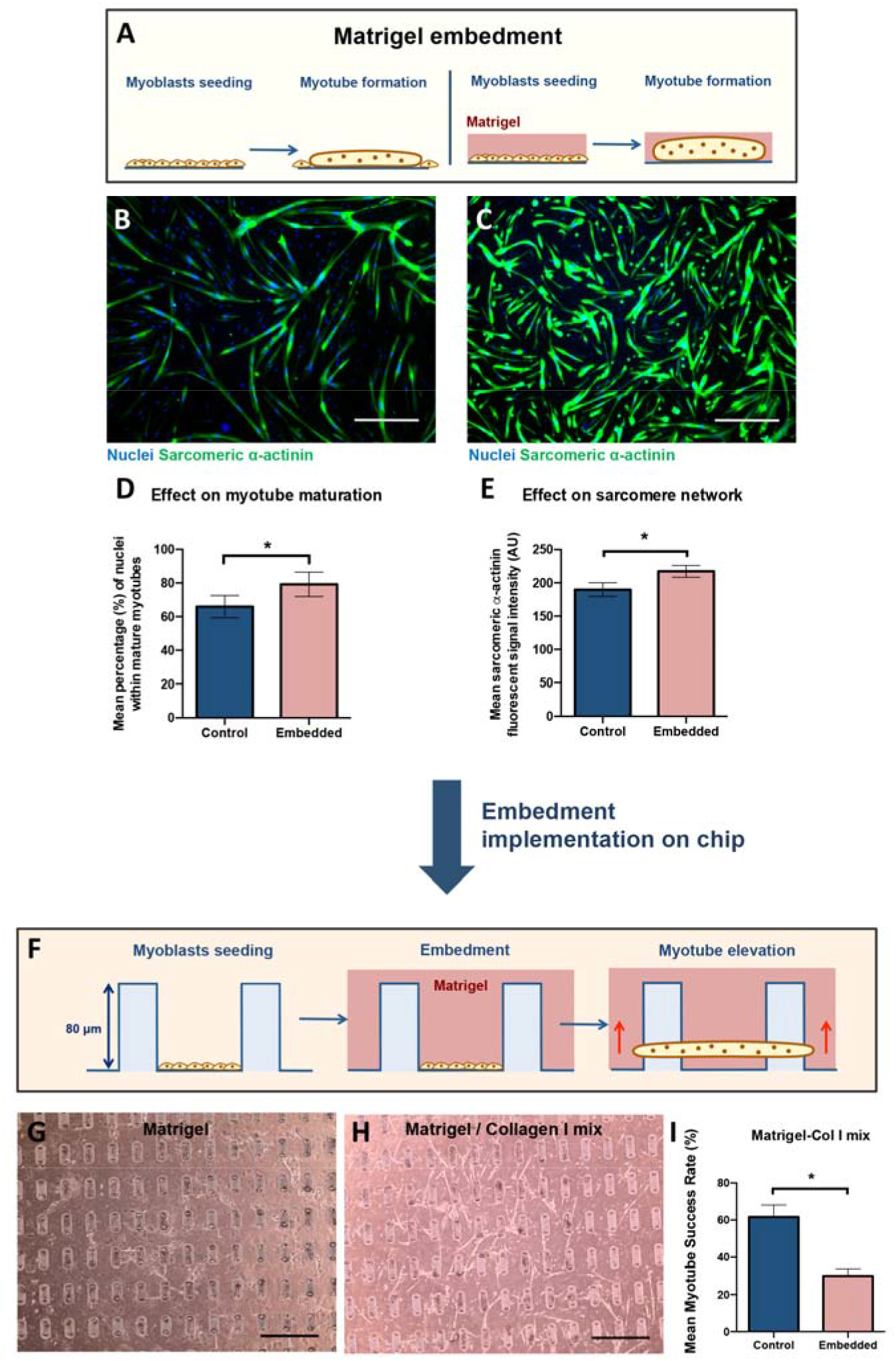
Incrementation of gel embedment methods in the system. **(A)** Schematic representation of Matrigel embedment protocol (right) in comparison to classic 2D culture of myotubes (left). **(B,C)** Representative images of immunofluorescent stained myotubes (from human immortalized myoblasts) after 6 days differentiation in normal 2D culture conditions (B) and Matrigel embedment conditions (C) (Sarcomeric *α*-actinin (green), nuclei (blue) scale bars = 500 μm). (D) Mean percentage ± s.e.m. of nuclei located within mature (Sarcomeric *α*-actinin-expressing) myotubes (from human immortalized (8220 and AB1190) myoblasts) after 6 days of differentiation with or without Matrigel embedment. *p<0.05 (Wilcoxon matched-pairs signed rank test), N=3. **(E)** Mean fluorescent signal intensity (Arbitrary Unit) ± s.e.m. from labeled Sarcomeric α-actinin of myotubes (from human immortalized (8220 and AB1190) myoblasts) after 6 days of differentiation with or without Matrigel embedment, *p<0.05 (Wilcoxon matched-pairs signed rank test), N=3. **(F)** Schematic representation of Matrigel embedment protocol adapted to 3D myotube culture on chip. **(G,H)** Transmitted light microscopic images of myotubes (from human immortalized (8220 and AB1190) myoblasts) after 7 days of differentiation with pure Matrigel (G), and Matrigel/Collagen I mix (H) embedments. Scale bars = 500 μm. I. Mean myotube culture success rate ± s.e.m. obtained from human immortalized (8220 and AB 1190) myoblasts after 7 days of differentiation on chip in normal culture conditions or Matrigel/Collagen I mix embedment. *p<0.05 (Mann-Whitney unpaired test), N=4.

**Fig. S3.**
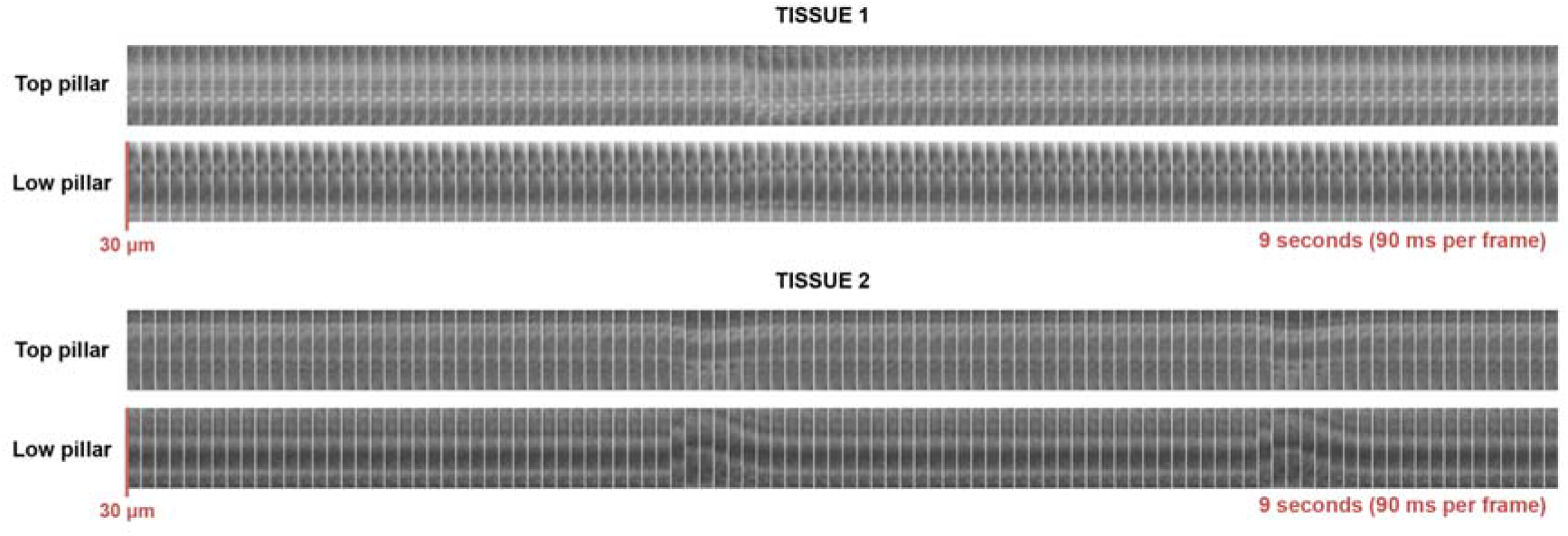
Kymographs. Transmitted light microscopic images of contracting 3D myotubes from human primary (AB678C53Q) myoblasts after 9 days of differentiation.

**Table S1.** Description table of cell lineages used in the study.

| Cell Line | Genotype | Phenotype | Lineage | Muscle | Sex | Age |
| --- | --- | --- | --- | --- | --- | --- |
| 8220 | - | Healthy | Immortalized | Paraspinal | Male | 12 yo |
| AB1190 | - | Healthy | Immortalized | Paravertebral | Male | 16 yo |
| EMD1365 | LMNA AAG, pLys32del c. del94-96 | LMNA related Congenital Muscular Dystrophy | Immortalized | Right gluteal | Male | 13 yo |
| AB678C53Q | - | Healthy | Primary | Quadriceps | Male | 53 yo |

**Table S2.** Description table of PDMS microfabricated pillar patterns used in the study.

| Pattern n° | Striation | Shape | Minor axis (μm) | Major axis (μm) | Axial ratio | Height (μm) |
| --- | --- | --- | --- | --- | --- | --- |
| 1 | 3.5 μm | Ellipse | 57 | 80.7 | 1.4 | 148.5 |
| 2 | 3.5 μm | Ellipse | 21.4 | 70.5 | 3.3 | 137 |
| 3 | 3.5 μm | round | 57 | - | 1 | 138 |
| 4 | 3.5 μm | Ellipse | 44 | 80 | 1.8 | 145 |
| 5 | 3.5 μm | Ellipse | 30 | 54 | 1.8 | 130 |
| 6 | 3.5 μm | Ellipse | 27 | 44.7 | 1.7 | 135 |
| 7 | None | Ellipse | 42.4 | 81.5 | 1.9 | 145 |
| 8 | None | Ellipse | 57 | - | 1 |  |

**Movie S1.**

**Spontaneous contractions of 3D myotubes obtained from primary human myoblasts (AB678C53Q) after 9 days of differentiation.**

**Movie S2.**

**Spontaneous contractions of 3D myotubes obtained from immortalized human myoblasts (8220) after 9 days of differentiation.**

**Movie S3.**

**Contractile activity of 3D myotubes obtained from L-CMD (EMD1365) patients immortalized myoblasts after 9 days of differentiation.**

## References

1. Gautel M, Djinović-Carugo K. The sarcomeric cytoskeleton: from molecules to motion. Lindstedt SL, Hoppeler HH, editors. Journal of Experimental Biology. 2016 Jan 1;219(2):135–45.

2. Powers JD, Malingen SA, Regnier M, Daniel TL. The Sliding Filament Theory Since Andrew Huxley: Multiscale and Multidisciplinary Muscle Research. Annu Rev Biophys. 2021 May 6;50(1):373–400.

3. Evano B, Tajbakhsh S. Skeletal muscle stem cells in comfort and stress. npj Regen Med. 2018 Dec;3(1):24.

4. Iaizzo PA, Durfee WK. Functional Force Assessment of Skeletal Muscles. In: Kramme R, Hoffmann K-P, Pozos RS, editors. Springer Handbook of Medical Technology.

5. Pimentel MR, Falcone S, Cadot B, Gomes ER. In Vitro Differentiation of Mature Myofibers for Live Imaging. JoVE. 2017 Jan 7;(119):55141.

6. Zhang H, Wen J, Bigot A, Chen J, Shang R, Mouly V, et al. Human myotube formation is determined by MyoD-Myomixer/Myomaker axis. Sci Adv. 2020 Dec;6(51):eabc4062.

7. Afshar Bakooshli M, Lippmann ES, Mulcahy B, Iyer N, Nguyen CT, Tung K, et al. A 3D culture model of innervated human skeletal muscle enables studies of the adult neuromuscular junction. eLife. 2019 May 14;8:e44530.

8. Kural MH, Billiar KL. Regulating tension in three-dimensional culture environments. Experimental Cell Research. 2013 Oct;319(16):2447–59.

9. Boudou T, Legant WR, Mu A, Borochin MA, Thavandiran N, Radisic M, et al. A Microfabricated Platform to Measure and Manipulate the Mechanics of Engineered Cardiac Microtissues. Tissue Engineering Part A. 2012 May;18(9–10):910–9.

10. Kalman B, Picart C, Boudou T. Quick and easy microfabrication of T-shaped cantilevers to generate arrays of microtissues. Biomed Microdevices. 2016 Jun;18(3):43.

11. Walker M, Godin M, Pelling AE. A vacuum-actuated microtissue stretcher for long-term exposure to oscillatory strain within a 3D matrix. Biomed Microdevices. 2018 Jun;20(2):43.

12. Legant WR, Pathak A, Yang MT, Deshpande VS, McMeeking RM, Chen CS. Microfabricated tissue gauges to measure and manipulate forces from 3D microtissues. Proc Natl Acad Sci U S A 2009 Jun 23;106(25):10097–102.

13. Kim D-H, Lipke EA, Kim P, Cheong R, Thompson S, Delannoy M, et al. Nanoscale cues regulate the structure and function of macroscopic cardiac tissue constructs. Proceedings of the National Academy of Sciences. 2010 Jan 12;107(2):565–70.

14. Gorji A, Toh PJY, Ong HT, Toh Y-C, Toyama Y, Kanchanawong P. Enhancement of Endothelialization by Topographical Features Is Mediated by PTP1B-Dependent Endothelial Adherens Junctions Remodeling. ACS Biomater Sci Eng. 2021 May 4.

15. Lam MT, Sim S, Zhu X, Takayama S. The effect of continuous wavy micropatterns on silicone substrates on the alignment of skeletal muscle myoblasts and myotubes. Biomaterials. 2006 Aug;27(24):4340–7.

16. Huang NF, Lee RJ, Li S. Engineering of aligned skeletal muscle by micropatterning. Am J Transl Res. 2010 Jan 1;2(1):43–55.

17. Théry M. Micropatterning as a tool to decipher cell morphogenesis and functions. Journal of Cell Science. 2010 Dec 15;123(24):4201–13.

18. Charest JL, García AJ, King WP. Myoblast alignment and differentiation on cell culture substrates with microscale topography and model chemistries. Biomaterials. 2007 May;28(13):2202–10.

19. Strale P-O, Azioune A, Bugnicourt G, Lecomte Y, Chahid M, Studer V. Multiprotein Printing by Light-Induced Molecular Adsorption. Adv Mater. 2016 Mar;28(10):2024–9.

20. Rao L, Qian Y, Khodabukus A, Ribar T, Bursac N. Engineering human pluripotent stem cells into a functional skeletal muscle tissue. Nat Commun. 2018 Dec;9(1):126.

21. Maffioletti SM, Sarcar S, Henderson ABH, Mannhardt I, Pinton L, Moyle LA, et al. Three-Dimensional Human iPSC-Derived Artificial Skeletal Muscles Model Muscular Dystrophies and Enable Multilineage Tissue Engineering. Cell Reports. 2018 Apr;23(3):899–908.

22. Sakar MS, Neal D, Boudou T, Borochin MA, Li Y, Weiss R, et al. Formation and optogenetic control of engineered 3D skeletal muscle bioactuators. Lab Chip. 2012 Dec 7;12(23):4976–85.

23. Cvetkovic C, Raman R, Chan V, Williams BJ, Tolish M, Bajaj P, et al. Three-dimensionally printed biological machines powered by skeletal muscle. Proceedings of the National Academy of Sciences. 2014 Jul 15;111(28):10125–30.

24. Mills RJ, Parker BL, Monnot P, Needham EJ, Vivien CJ, Ferguson C, et al. Development of a human skeletal micro muscle platform with pacing capabilities. Biomaterials. 2019;198:217–27.

25. Osaki T, Uzel SGM, Kamm RD. On-chip 3D neuromuscular model for drug screening and precision medicine in neuromuscular disease. Nat Protoc. 2020 Feb;15(2):421–49.

26. Saez A, Buguin A, Silberzan P, Ladoux B. Is the mechanical activity of epithelial cells controlled by deformations or forces? Biophys J. 2005 Dec;89(6):L52–54.

27. Afshar ME, Abraha HY, Bakooshli MA, Davoudi S, Thavandiran N, Tung K, et al. A 96-well culture platform enables longitudinal analyses of engineered human skeletal muscle microtissue strength. Sci Rep. 2020 Dec;10(1):6918.

28. Bertrand AT, Ziaei S, Ehret C, Duchemin H, Mamchaoui K, Bigot A, et al. Cellular microenvironments reveal defective mechanosensing responses and elevated YAP signaling in LMNA-mutated muscle precursors. Journal of Cell Science. 2014 Jul 1;127(13):2873–84.

29. Bruusgaard JC, Johansen IB, Egner IM, Rana ZA, Gundersen K. Myonuclei acquired by overload exercise precede hypertrophy and are not lost on detraining. Proceedings of the National Academy of Sciences. 2010 Aug 24;107(34):15111–6.

30. Hastings RL, Massopust RT, Haddix SG, Lee Y il, Thompson WJ. Exclusive vital labeling of myonuclei for studying myonuclear arrangement in mouse skeletal muscle tissue. Skeletal Muscle. 2020 Dec;10(1):15.

31. Sullivan T, Escalante-Alcalde D, Bhatt H, Anver M, Bhat N, Nagashima K, et al. Loss of a-Type Lamin Expression Compromises Nuclear Envelope Integrity Leading to Muscular Dystrophy. Journal of Cell Biology. 1999 Nov 29;147(5):913–20.

32. Bertrand AT, Brull A, Azibani F, Benarroch L, Chikhaoui K, Stewart CL, et al. Lamin A/C Assembly Defects in LMNA-Congenital Muscular Dystrophy Is Responsible for the Increased Severity of the Disease Compared with Emery-Dreifiiss Muscular Dystrophy. Cells. 2020 Mar 31;9(4):844.

33. Earle AJ, Kirby TJ, Fedorchak GR, Isermann P, Patel J, Iruvanti S, et al. Mutant lamins cause nuclear envelope rupture and DNA damage in skeletal muscle cells. Nat Mater. 2020;19(4):464–73.

34. Saez A, Ghibaudo M, Buguin A, Silberzan P, Ladoux B. Rigidity-driven growth and migration of epithelial cells on microstructured anisotropic substrates. Proceedings of the National Academy of Sciences. 2007 May 15;104(20):8281–6.

